# *Diffractograph:* single-particle reconstruction using electron-diffraction data

**DOI:** 10.64898/2026.10.14.759276

**Authors:** Senik Matinyan, Jan Pieter Abrahams

## Abstract

We recently showed that narrow-beam electron diffraction of single particles provides higher signal-to-noise ratios and lower radiation damage than imaging. Here we introduce *Diffractograph*, a suite for reconstructing three-dimensional structure from these data.

*Diffractograph* selects mutually compatible diffraction frames, groups them into one or more models and discards incompatible frames. In the *ab initio* workflow, this is done without prior structural information. In the reference-guided workflow, a low-resolution electrostatic potential map from single-particle cryo-EM is used to select compatible frames and determine their orientations. The oriented frames are combined into a three-dimensional intensity distribution. The reference map supplies low-resolution phases for the oversampled structure factors, and density modification extends these phases to higher resolution. For synthetic 200-keV diffraction data extending to 2.2 Å, *Diffractograph* retained all frames and recovered the correct orientation of every retained frame with a median plane-normal error of 0.27°. These results establish a computational workflow for structure determination using single-particle electron-diffraction data.

## 1. Introduction

Our recent 4D-para-STEM study showed that reducing the illuminated area reduced observed radiation damage. Narrow beam diffraction also gave higher reciprocal-space signal-to-noise ratios than conventional cryo-EM imaging of thin, non-crystalline specimens [1]. Narrow-beam diffraction may therefore extract more structural information from a radiation-limited specimen. It also produces many phase-free patterns with unknown orientations. Some frames may contain only solvent or support scattering; others may represent different structures. Machine learning can help identifying useful hits [2], but those hits must still be oriented in three dimensions. *Diffractograph* addresses the latter problem and allows self-consistent models to emerge.

The geometry of orientation recovery is simple. The Fourier transform of a projection forms a central plane through the three-dimensional transform. Two such planes intersect along a common line, and the three common lines among three projections determine their relative orientations up to handedness [3]. Noise makes the correct line difficult to distinguish from plausible alternatives. Voting, spectral synchronization and statistical weighting use agreement among many images to resolve this uncertainty [4,5]. These methods were developed for projection images, whose complex Fourier transforms retain phase. Diffraction records only intensity. Diffraction intensities contain no measured phases, but they are invariant to translation. This removes the two positional parameters required when aligning projection images.

Common intersections have also been used directly with phase-free diffraction data: common lines for flat-Ewald XFEL patterns [6], common arcs for curved Ewald spheres [7,8], and multiple common lines for weak patterns [9]. EMC instead alternates between assigning orientations and refining a three-dimensional intensity model [10]. Most of this work concerns XFEL data, where radiation damage and experimental background remain major constraints [11]. Electrons provide more useful elastic scattering per unit radiation damage [12], while narrow-beam illumination reduces solvent background, beam-induced motion and observed diffraction decay [1].

The short electron wavelength makes the measured Ewald surface nearly flat. Under the kinematical approximation, Friedel symmetry therefore makes each pattern approximately C2 symmetric, leaving a 180° ambiguity about the incident beam. Dynamical scattering breaks Friedel symmetry and has been used to determine absolute structure in crystals [13]. We asked whether this weak asymmetry could also resolve the relative binary orientations of continuous single-particle patterns. If so, each pattern could be assigned to the appropriate Ewald surface and the patterns assembled in three dimensions. Reversing every assignment would still give the opposite hand.

*Diffractograph* avoids selecting a single common line for each pair. Instead, it retains the probability of the possible relative rotations and common-line directions, including the possibility that the apparent match is uninformative. These probabilities form the edges of a graph, calculated only where needed. Consistent edges establish small orientation models, which are then merged and refined. Incompatible models remain separate, and unsupported frames need not be assigned. This *ab initio* procedure requires no prior structural information. Alternatively, a low-resolution map from single-particle cryo-EM can be Fourier transformed and used to select compatible frames and guide their orientations. Once the centrosymmetric orientations are known, the dynamical-scattering signal resolves the remaining relative C1 orientations and the patterns are assembled on curved Ewald surfaces.

The reconstructed intensity distribution supplies structure-factor amplitudes but not phases. Oversampling provides constraints for phase recovery but does not by itself determine a unique solution [14]. In the reference-guided workflow, the low-resolution map provides initial phases by molecular replacement. Density modification then uses the reconstructed oversampled amplitudes to extend these phases to higher resolution. We validate the orientation procedure using 1,741 synthetic electron-diffraction patterns with known orientations, Ewald-sphere curvature and dynamical scattering.

## 2. Algorithm

### 2.1 C2-anisotropy analysis

First, the beam centre of each diffraction frame is determined from its C2 symmetry. Solvent and support scattering are predominantly isotropic, whereas particle diffraction is generally anisotropic. Under the kinematical, flat-Ewald approximation, this variation obeys Friedel symmetry: intensities at opposite points on a resolution ring are equal.

*Diffractograph* uses this symmetry to estimate and then refine the beam centre of each frame. It assigns a C2-anisotropy evidence score between zero and one. For two opposite pixels, *a* and *b*, it calculates the Friedel-even and Friedel-odd components

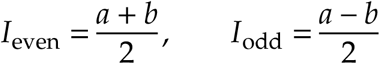

The isotropic radial mean is subtracted from the even component. The even and odd powers (*P*_even/odd_) are then accumulated over the selected resolution range, with weights for ring radius and valid detector coverage. The evidence score is

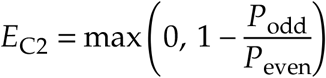

Unstructured counting noise contributes equally to the even and odd powers, giving a score near zero. C2-symmetric particle diffraction adds predominantly to the even power, raising the score towards one. The odd power can also contain beam-centre errors, detector artefacts, structured background, Ewald-sphere curvature and dynamical scattering. The score therefore ranks evidence for useful C2-symmetric anisotropy; it is not a calibrated probability that a frame contains a particle. The odd signal is not removed from the underlying data and remains available for subsequent C1 analysis.

The survey records the input filenames, refined beam centres, C2-anisotropy evidence scores and experimental geometry in the workflow JSON output. Frames can be filtered by a minimum C2-anisotropy evidence score, by retaining a chosen highest-scoring fraction, or by both criteria. This removes evidently solvent-dominated frames before the more expensive polar resampling and pairwise-intersection calculations.

### 2.2 Polar resampling and anisotropy frames

Each selected frame is resampled on polar rings about its beam centre (Fig. 1; all figures use the 1AUP dataset used in *Results*). By default, the radial spacing is one detector pixel. The number of angular samples increases with radius so that their spacing along each ring also remains close to one pixel. Ring sizes are adjusted for efficient Fourier transformation.

**Figure 1.**
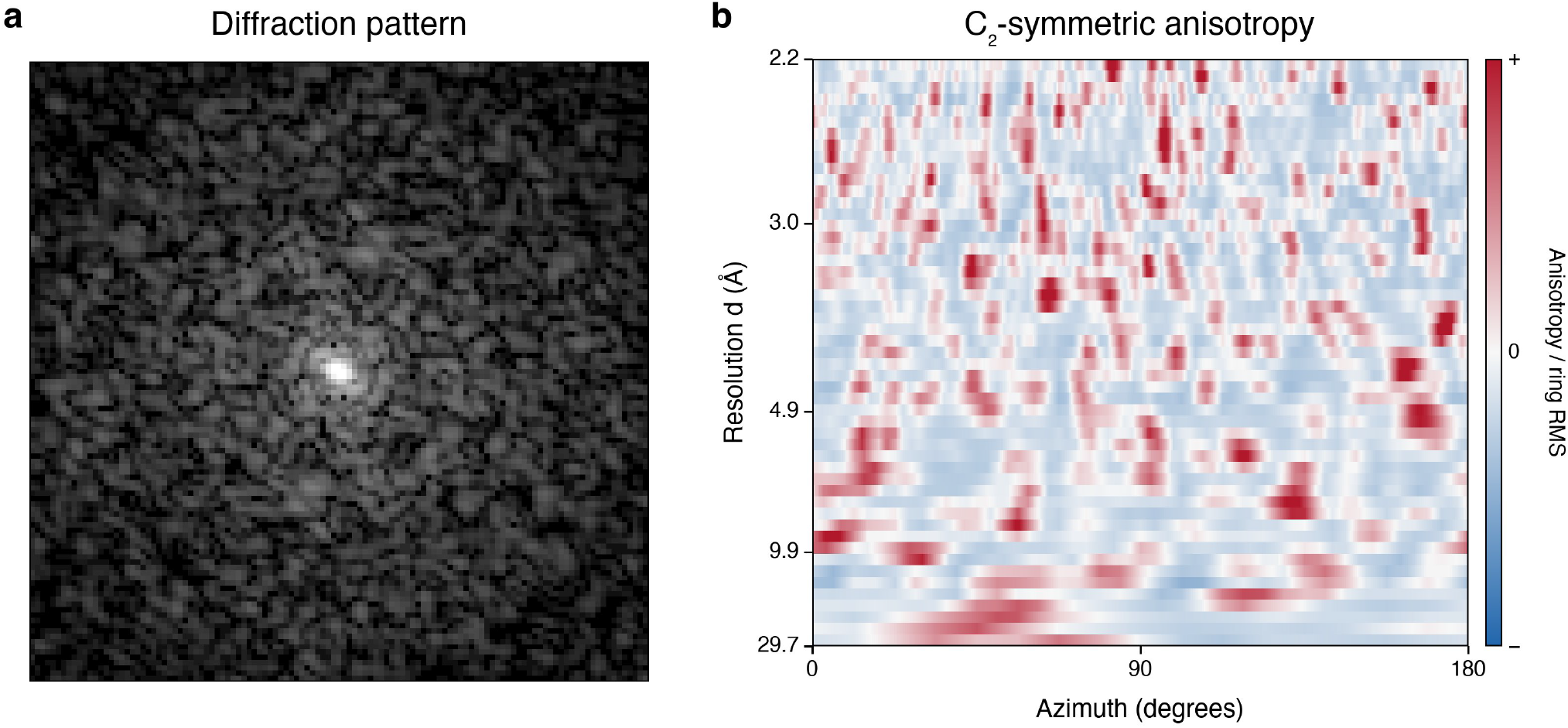
Conversion of a diffraction pattern into its anisotropy representation. **a**, Representative synthetic continuous electron-diffraction pattern. The complete measured intensity range I **x** of 0–65,535 counts is shown using log 1 + I **x** scaling without intensity clipping. Five overloaded pixels surrounding the direct beam were masked. **b**, Corresponding polar anisotropy representation after centrosymmetric averaging and subtraction of the isotropic intensity on each radial ring. The azimuthal domain is reduced to p radians by the C2 symmetry. For visualization, each ring was divided by its root-mean-square anisotropy, and the diverging colour scale was limited symmetrically at the 99.5th percentile of the absolute normalized values. Red and blue indicate intensities above and below the radial mean, respectively.

Samples separated by π are averaged, producing a C2-symmetric ring with angular period π. If only one member of a Friedel pair is valid, its value is retained; if neither is valid, the sample remains missing. The radial mean is then subtracted from each ring and stored separately. The remaining variation contains the angular diffraction signal from the particle, possible particle–environment or structured-background contributions and counting noise. Detector masks and incomplete angular coverage are recorded as missing observations and valid-coverage fractions. Ring powers and angular Fourier transforms are cached for normalizing ring correlations. We call the resulting C2-symmetric object an *anisotropy frame*, and refer to it as *frame* in the remainder of the text. An anisotropy frame is not an image or volume, but a compact, flat-Ewald representation of the orientational information in one diffraction pattern.

### 2.3. Pairwise intersections are probabilistic graph edges

For two anisotropy frames, *Diffractograph* searches for the in-plane rotation, α, that rotates the second frame onto the first, and the common-line direction in the first frame, γ. In the second frame the common line direction is α + γ.

The search proceeds in stages. Fourier transforms are first used to correlate the radial intensity variation of corresponding rings. Each ring correlation is normalized by its power, weighted by the fraction of valid detector pixels and added to overlapping, sector-specific rotation correlation profiles. The calculation is repeated with the order of the two frames reversed. Candidate rotations are favoured when the forward and reverse searches agree, and up to a configurable number of distinct candidates are retained (six by default). A local Gaussian fit estimates the position and width of each rotation maximum.

For each candidate rotation, *Diffractograph* searches the compatible common-line directions. In each direction γ, the intensity variation in the first frame is compared with that at γ + α in the second. Values on each ring are divided by the square root of the corresponding ring power, so the comparison depends on the agreement of the radial profiles rather than their absolute scale. The best-supported positive correlation supplies the common-line direction. A conservative fit to the peak gives its variance.

Correlations are then recalculated at several nearby rotations. A quadratic fit to the logarithm of the positive correlations estimates the coupling between errors in α and γ. This cross term is combined with the two independently estimated variances to give the covariance matrix of a wrapped bivariate Gaussian. A complete two-dimensional correlation surface is therefore neither calculated nor stored.

A sharp maximum does not guarantee that the proposed intersection is correct. The support for competing candidates is used to assign a uniform background component:

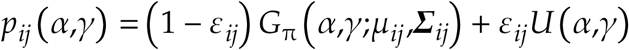

Here, *G*_π_ is a bivariate Gaussian wrapped with period π in both angular coordinates, *µ*_*ij*_ is its maximum and Σ_*ij*_ its covariance. *U* is uniform over the same angular domain. The mixture weight *ε*_*ij*_ represents support for an uninformative or incorrect match; it should not be interpreted as a calibrated posterior probability. If no valid intersection is found, the edge is entirely uniform.

Figure 2 illustrates how the information in an edge can change with relative tilt. For nearly coincident planes, the relative rotation may be well determined while the common-line direction remains broad. At larger tilts, the common-line direction can become more precise. A narrow Gaussian does not necessarily imply a reliable match: competing maxima can still give a large uniform component. The wrapped Gaussian mixture can be evaluated at arbitrary angles and has analytic first and second derivatives. This permits efficient refinement without interpolation of a sampled correlation surface.

**Figure 2.**
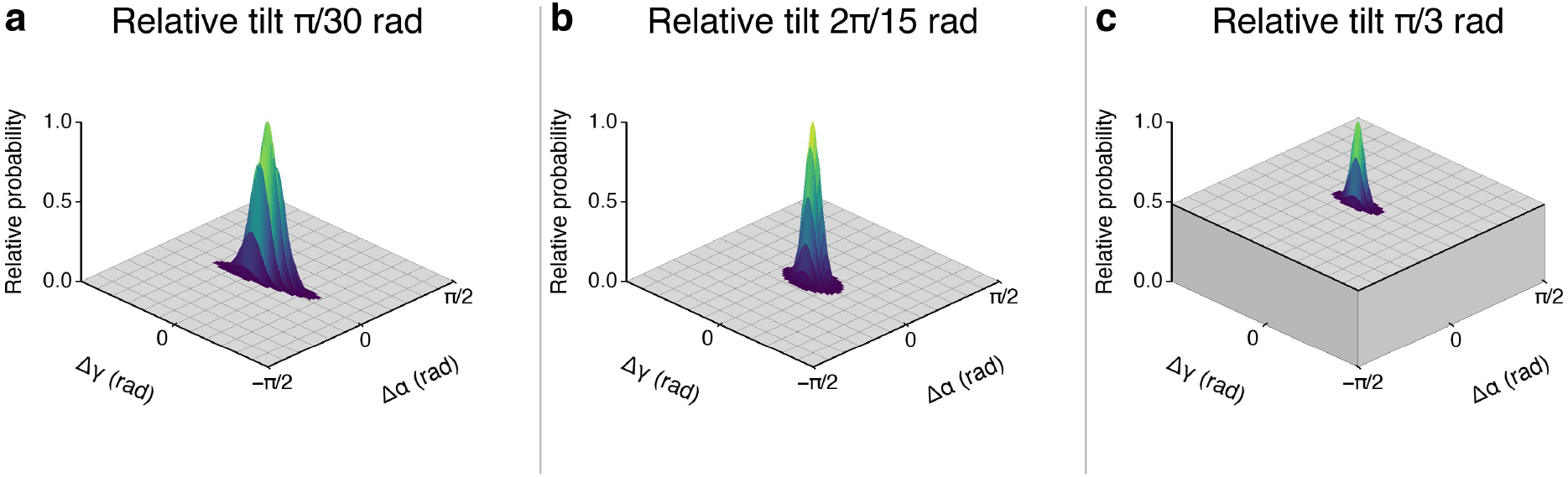
Pairwise intersection evidence changes with relative tilt. Periodic joint probability distributions for relative in-plane rotation, Δα, and common-line direction, Δγ, at relative tilts of (a) π/30, (b) 2π/15 and (c) π/3 rad. The fitted standard deviations σ_a_, σ_g_ and correlation coefficients ρ are respectively (0.044, 0.210, −0.610), (0.063, 0.119, −0.052) and (0.034, 0.107, 0.001), in radians. In panel c, the grey pedestal represents the uniform mixture weight of 0.486; the remaining 0.514 is assigned to the localized Gaussian hypothesis. Surfaces were centred on their fitted maxima and normalized to a peak height of one.

Frames are the nodes of a graph. Each calculated pairwise intersection is an edge whose probability distribution describes the compatible relative rotation and common-line direction. Edges are calculated only when needed and then cached, avoiding construction of the complete pairwise graph. If every pair in an *N*-frame model were evaluated, the graph would provide *N N* − 1 angular observations for 3*N* − 3 orientation parameters, a nominal ratio of *N*/3. The calculated graph is sparse for larger models, so this fully connected-graph count describes only how the number of potential pairwise constraints scales with model size.

### 2.4. Ab initio orientation

#### 2.4.1 Local compatibility seeds three-dimensional models

A single graph edge relates two diffraction planes but does not establish a three-dimensional coordinate system. Diffractograph therefore searches overlapping graph neighbourhoods for sets of three mutually connected frames, here called triplets. As a rapid initial filter, it selects triplets for which the product of the three edge signal probabilities exceeds a configurable threshold. For each frame, only the most strongly supported triplets are carried forward. If their three pairwise intersections are mutually consistent, they determine the relative orientations of the three diffraction planes, apart from an overall rotation and hand.

These locally consistent triplets are the initial orientation models: sets of frames with mutually consistent orientation matrices. Regardless of its size, an orientation model assigns a proper 3 × 3 rotation matrix *R*_*i*_ to each of its frames. Its quality is the mean log-likelihood ratio over its supporting edges:

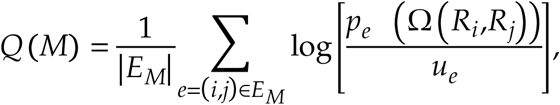

Here, *E*_*M*_ is the set of supporting edges, *p*_*e*_ is the probability density of edge *e* (Fig. 2), and Ω(*R*_*i*_,*R*_*j*_) gives the relative rotation and common-line direction inferred from *R*_*i*_ and *R*_*j*_. The uniform density *u*_*e*_ represents an edge with no angular information. Because it is independent of the fitted orientations, it does not affect their optimum. Dividing by the number of edges makes model quality comparable across model sizes; *Q(M)* > 0 means that the fitted geometry is favoured over an uninformative match on average.

Once an orientation model contains redundant supporting edges, its rotation matrices can be improved by optimizing *Q(M)*. Current frame rotations imply a rotation and common-line direction for each supporting edge. The fit uses the full two-dimensional probability density at those predicted angles (Fig. 2), rather than only the location of its maximum. An edge with a broad peak, or with correlated rotation and common-line angles, then constrains only the combinations of orientation changes that it determines reliably. When an edge is noisy and has a large uniform component (Fig. 2c), its probability density changes less with the predicted angles and therefore has less influence on the fit.

Triplets sharing an edge are combined into candidate quartets. Any missing pairwise intersections are calculated, and the four orientations are refined jointly against all six edge distributions. Refined quartets sharing three frames are then merged into quintets. Missing intersections are again calculated, and the five orientations are refined together. A quintet is retained only when its fitted geometry is supported more strongly than an uninformative match. These locally consistent quintets seed the subsequent model consolidation.

For initial seeding, the frames are placed in a reproducible pseudorandom circular order. Each frame is connected to a user-specified number of nearby frames in that ordering, producing overlapping local neighbourhoods. Required intersections are calculated as the neighbourhoods are traversed, and qualifying triplets, quartets and quintets are generated and refined. Because adjacent neighbourhoods overlap, frames can occur in several seed orientation models.

#### 2.4.2. Model merging converts local evidence into a global reconstruction

The refined quintets provide many overlapping local orientation models. *Diffractograph* merges models that share sufficient frames to relate their coordinate systems. Small models require at least four shared frames. When both models contain more than 32 frames, the required overlap increases with the size of the smaller model, from at least 10 shared frames to a maximum of 16.

The shared frames are used to test the relative hand and to align the coordinate systems of the two parent models. For large overlaps, the alignment is fitted jointly over the shared orientations. For smaller overlaps, alternative alignments generated from the shared frames and their C2-equivalent orientations are tested. The alignment that gives the best joint likelihood initializes the union.

The frames are placed in a common coordinate system. For unions of up to 32 frames, all missing pairwise intersections are calculated. Larger unions retain their existing graph edges and calculate selected interface edges that connect the two parent models. The union is then refined against its supporting edge distributions.

Before a candidate union is accepted, additional locally compatible edges are calculated where the fitted plane normals are within a specified tilt limit. Frames with unusually poor residuals are refitted and, if they remain inconsistent, rejected. The resulting model is accepted only when its mean edge log-likelihood ratio is non-negative. Accepted unions replace their parent models, and models wholly contained within a larger model are removed.

Consolidation proceeds in rounds until no further supported unions remain. The final models may represent separate particle populations, distinct orientation groups, or unsupported frames that could not be assigned consistently.

### 2.5. Reference-based orientation

A three-dimensional reference gives *Diffractograph* a starting point for orienting diffraction frames. It allows each frame to be rotated in three dimensions relative to something already known. This orientation is provisional: intersections between experimental frames are then used to test and refine it.

The reference can take three forms. Either a low-resolution cryo-EM map or a three-dimensional diffraction intensity volume produced by an earlier *Diffractograph* run provides a three-dimensional diffractogram against which frame orientations can be fitted. Alternatively, an existing *Diffractograph* orientation model in JSON format provides experimental frames whose orientations are already known.

Frames with positive experimental support are eligible for addition. They are ranked by how well their intensities correlate with nearby model frames along the common lines predicted by their provisional orientations. This lets the best-matching frames strengthen the model first.

Frames are added in small groups to the growing model. For each group, *Diffractograph* makes a trial model and refines the orientations of the existing and new frames together. It accepts the group only if the original frames remain in the model and their fit does not worsen. If the group fails, it is split and its frames are tried in smaller groups. After the model has grown, frames that were not accepted are tried again; this continues until no further frames are added.

### 2.6 Orientation refinement and model completion

Orientation refinement improves an existing orientation model using the experimental intersections alone. *Diffractograph* first selects many small, overlapping seed models from the input model. Their frames are joined by significant, informative edges, and each seed contains substantially more independent pairwise observations than orientation parameters. The seed models are therefore well overdetermined, while remaining small enough that their orientations can move efficiently away from suboptimal values imposed by a larger input model. Each seed is refined against its pairwise intersections. The refined seeds are then consolidated using the same merging procedure as in the *ab initio* workflow. As models grow, their orientations are refined jointly and poorly supported frames can be refitted or rejected. Repeating this procedure with different overlapping seeds allows local orientation errors in the input model to be corrected without requiring an external reference.

An orientation model can be completed by providing additional frames, which will be added if they fit sufficiently well.

### 2.7. Dynamical scattering resolves Ewald sphere assignments

The C2 orientation of a frame leaves two possible C1 orientations. They differ by a rotation of π about the incident beam and place the frame on opposite faces of the Ewald sphere. The Friedel-odd intensity allows distinguishes these alternatives

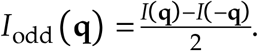

In the kinematical approximation and ignoring Ewald sphere curvature, Friedel symmetry holds, and *I*_odd_ (**q**) is zero. However, dynamical scattering breaks Friedel symmetry at low resolution, resulting in a small non-zero *I*_odd_ (**q**). Assigning a frame to its alternative C1 orientation reverses the sign of *I*_odd_ (**q**). Two frames with close orientations and matching C1 assignments should therefore have positively correlated *I*_odd_ (**q**) values along the common line, while opposite assignments are negative correlated.

The frames are arranged into overlapping local groups with close orientations. All pairwise odd-component correlations are evaluated within each group, considering both the C2 orientation and after a rotation of π. Within each group, all permutations are considered and the combination with the largest summed pair correlation is selected. The overlapping groups determine one another’s signs through the frames they share. Groups with the strongest overlap are mutually aligned first. For each frame, its prediction is calculated from all other frames. A negative correlation between the observed and predicted odd signal rotates it by π.

### 2.8. Three-dimensional reciprocal-space reconstruction

#### 2.8.1 Reciprocal-space slice insertion

After the C1 orientations have been assigned, *Diffractograph* uses the radial mean-subtracted intensity samples from the corresponding detector frames. A frame-specific overall scale and radial offset are applied as described in Section 2.8.2. The corrected pixel value is the anisotropic signal inserted into the three-dimensional volume.

Each pixel’s scattering vector is calculated from the calibrated detector geometry on the electron Ewald sphere, then rotated into the common reciprocal-space coordinate system using the frame’s C1 orientation. This inserts the anisotropic pixel samples at their curved-Ewald coordinates, rather than gridding the flat polar anisotropy arrays used for orientation.

Each anisotropic pixel sample contributes to neighbouring voxels by trilinear interpolation. *Diffractograph* accumulates weighted intensity and interpolation weight separately; their ratio gives the reconstructed intensity at each observed voxel. Voxels with zero sampling weight remain unobserved. Insertion may be extended along the rotated detector-plane normal with a Gaussian kernel whose width is estimated from the low-tilt reciprocal-intensity correlation. The intensity and sampling-weight fields are written as MRC files. When MTZ output is requested, *Diffractograph* also writes P1 reflection data, including intensities and their standard uncertainties, in a cubic unit cell set by the reciprocal-grid spacing.

#### 2.8.2. Frame scales, radial offsets and Wilson pedestal

The initial 3D volume is built from the radial-mean-subtracted pixel samples using initial frame scales stored in the orientation model. Now, the radial average of just the protein component needs to be added back in. These frame-specific radial offsets start at zero. *Diffractograph* then alternates between fitting each frame against a leave-one-out prediction from the current volume and updating the scale and radial offsets of the mean-subtracted frames.

At each iteration, *Diffractograph* calculates the intensities that each frame is expected to have, based on the values of all other frames in the slice defined by its orientation. It compares the prediction with the frame’s measured intensities and adjusts two things: one scale for the whole frame, and an additive offset for each resolution ring. The scale corrects differences in overall contrast; the offsets correct differences in ring means.

The frame scales are normalized so their geometric mean is one. At each ring, the offsets are centered so their sample-weighted mean is zero. This keeps the corrections from changing the overall scale or adding a shared radial profile. The program applies half of each fitted correction, updates the volume, and repeats. It gives less weight to uncertain offsets and smooths them between eligible neighboring rings.

For a pixel in frame *i*, the corrected intensity is

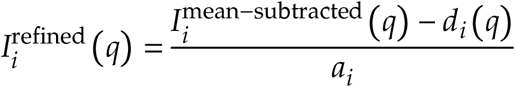

where *a*_*i*_ is the relative frame scale and *d*_*i*_ (*q*) is its interpolated radial offset.

After this refinement, the shared Wilson pedestal is calculated from the reconstructed volume before pedestal addition. For each resolution shell, let 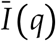 be its mean intensity and σ_*I*_ (*q*) its RMS intensity about that mean. The pedestal profile is

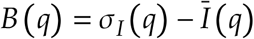

For an exponential Wilson intensity distribution, the mean and standard deviation are equal. Adding *B (q)* therefore raises the shell mean toward its RMS. The global Wilson profile offset is interpolated linearly in reciprocal frequency and added to supported voxels. The resulting intensities are on a common scale and have their protein-like isotropic pedestal restored (fig. 3).

**Figure 3.**
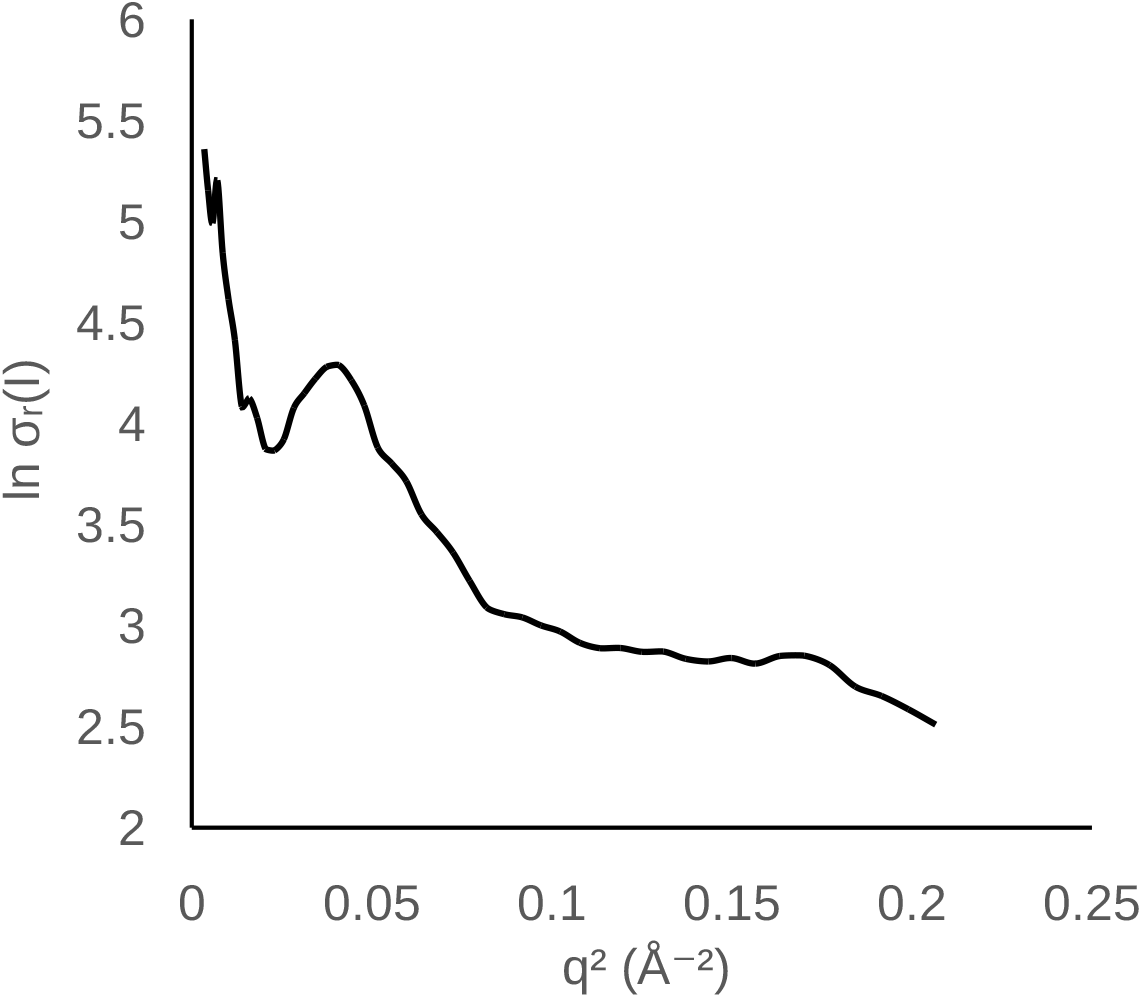
Wilson plot of the reconstructed intensity. The curve shows the shell mean intensity, which coincides with the RMS intensity about the mean after the global pedestal is added, as expected for Wilson-distributed intensities. The maximum around q^2^ ≈ 0.04 is typical for protein and is caused by its secondary structure.

### 2.9. Phasing by molecular replacement and density modification

The reconstructed intensity volume provides oversampled structure-factor amplitudes but not phases. For use with established molecular-replacement programs, the intensities and their standard deviations can be written to an MTZ file in CCP4 format. When no atomic model is available, *Diffractograph* can instead use a low-resolution cryo-EM map to start phase extension, followed by density modification with the *Solomon* solvent-flipping algorithm [16, 17].

Before estimating amplitudes, *Diffractograph* merges measured Friedel mates and subtracts a scaled Wilson pedestal from the intensities. The pedestal profile comes from the 3D reconstruction; its scale defaults to 0.15. This is not a standard procedure in density modification, but we found that for oversampled data it was crucial for success. This subtraction reduces the radial background before negative intensities are handled. Observations more than four standard deviations below zero are rejected; weaker negative observations are retained and converted to amplitudes with the French–Wilson procedure [18], which accounts for measurement uncertainty and a non-negative intensity prior.

To establish the initial phase prior, *Diffractograph* Fourier transforms the cryo-EM map and aligns and scales its calculated intensities against the experimental intensities over their shared resolution range. At each resolution shell, the agreement between map and experiment determines how strongly the map phases are σ_*A*_-weighted, accounting for experimental uncertainty [19]. Map phases provide the initial phases within the map’s resolution range.

Phase extension proceeds by opening progressively higher-resolution shells. New coefficients are seeded from the current solvent-modified map. At each cycle, the coefficients are transformed to real space, density modification is applied using a molecular support, and the result is transformed back to reciprocal space. The measured amplitudes constrain observed coefficients, with their influence weighted by uncertainty; the map-derived phase prior also contributes where available. Once the active outer shell has converged, the next shell is opened.

The user may supply the molecular support, or *Diffractograph* can estimate it from the low-resolution map and molecular weight. It selects a molecular core from the map and expands that core to form a solvent mask. As resolution increases, the support is updated from the density and the mask becomes more closely defined. In the solvent region, the *Solomon* procedure flips the density about the solvent level; density within the molecular support is retained. Repeated density modification, amplitude constraints and phase-prior weighting extend the phases beyond the cryo-EM map’s resolution.

## 3. Results

We used 1,741 synthetic, continuous electron-diffraction patterns (117×117 pixels) with known orientations. These were generated by AbTEM [20] of a monomer of glutamate dehydrogenase (pdb code 1AUP). The 200-keV simulation extended to 1.9 Å and included Ewald-sphere curvature and dynamical scattering (see Methods). Redundant zero-tilt patterns were removed. Their known orientations were used only for validation.

*Ab initio* orientation produced a model containing 1,739 oriented frames with median and 90th-percentile plane-normal errors of 1.25° and 4.02°, respectively. On an Apple Mac Studio (M1 Ultra), this took about 4:40 min. Two rounds of refinement (5:40 min), improved the orientation angles to 0.33° and 0.87°. The model was completed in 2 s to include all 1741 frames. Subsequent C1 assignment rotated all 1741 frames in the correct orientation, ensuring proper Ewald sphere alignment (4 s).

Orientation using the 1AUP density map as a reference in the resolution range between 10 and 30 Å, and using frame resolutions between 10 and 3 Å, produced a 1739-frame orientation model with median and 90th-percentile plane-normal errors of 1.40° and 3.52°, respectively (36 s). Three rounds of refinement improved these angles to 0.28° and 0.76° (9:40 min).

Three-dimensional intensity reconstruction (2:10 min) produced good leave-one-out agreement between each frame’s measured intensities and the intensities predicted from the 3D volume reconstructed without that frame, as assessed two-dimensional FSC (Fig. 4).

**Figure 4.**
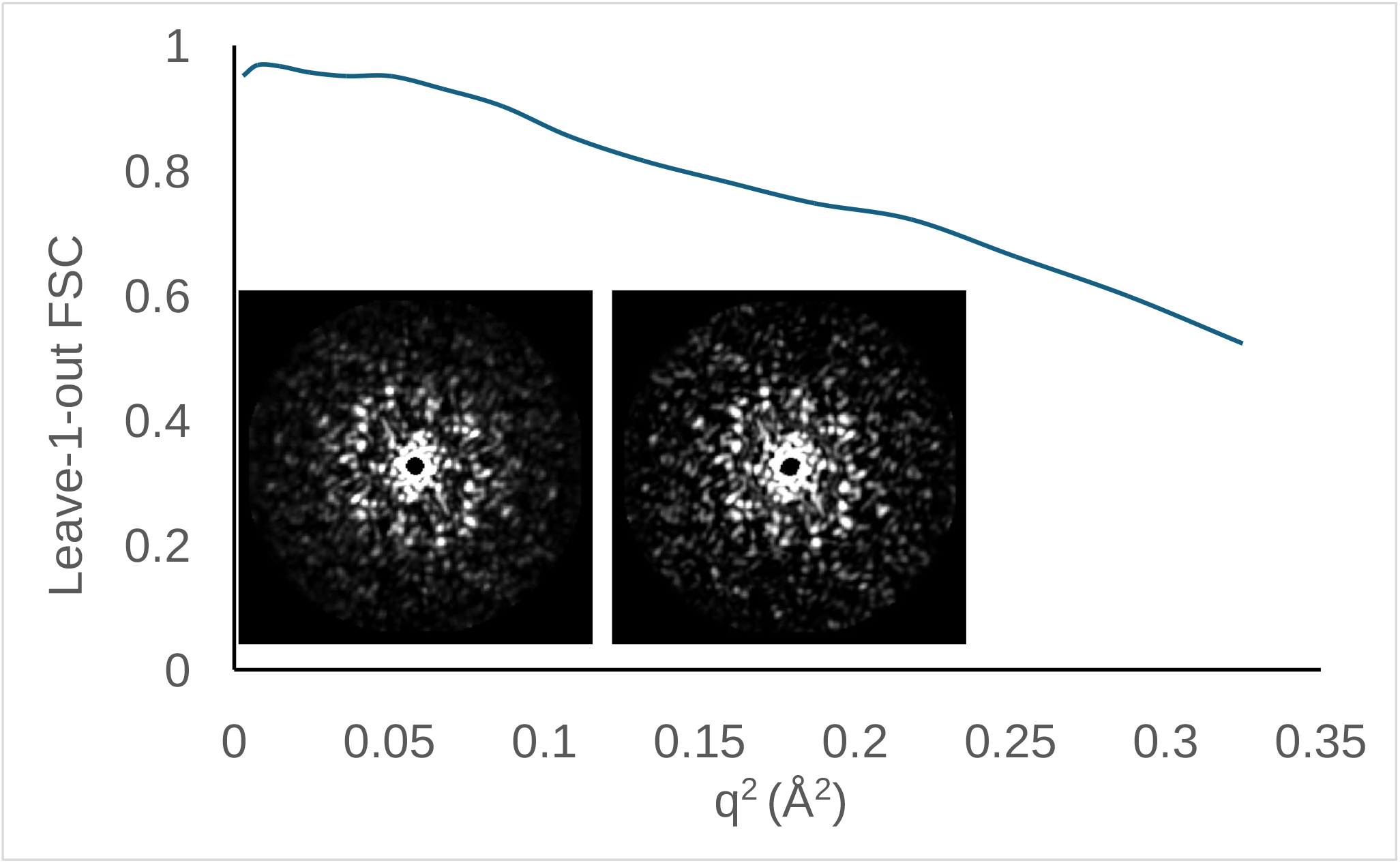
Three-dimensional reciprocal space intensity reconstruction. A 250 × 250×250 three-dimensional map with structure factors was assessed by the mean leave-one-out FSC statistic, described in the text. It provides the average Fourier shell correlation as a function of resolution of fits of each frame with the combined value of all other frames. Its high values (higher than 0.9 for resolutions between 30 and 3.5 Å, decreasing to 0.66 at 1.9 Å) indicate reconstruction was successful. The insert shows a single frame (right) and a slice through reciprocal space, computed using all other intensities in the dataset.

**Figure 5.**
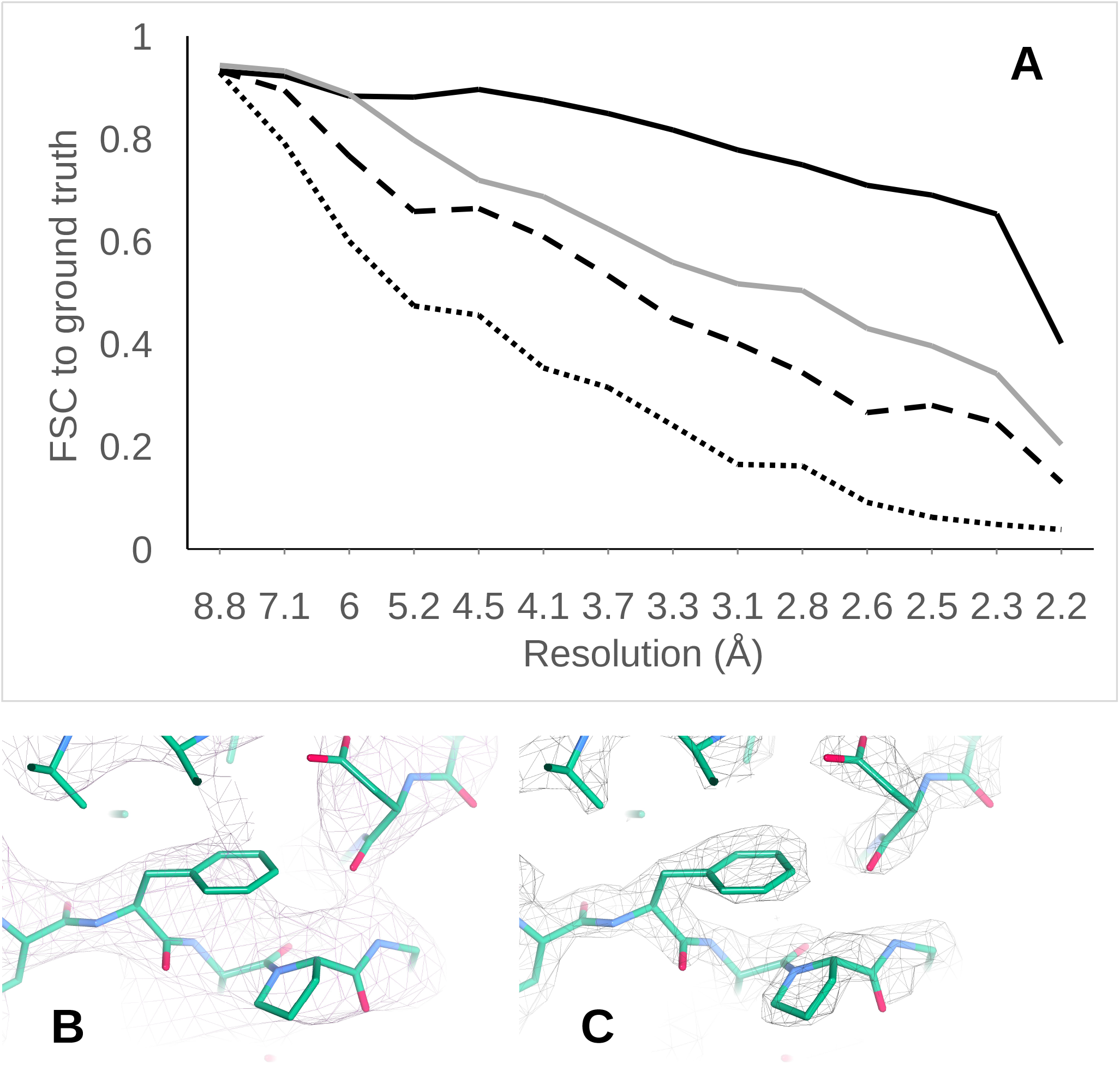
Phasing and phase extension of reconstructed three-dimensional intensity data. (A) FSC curves of phase extensions using molecular replacement density maps of different resolution: 8 Å (dotted line), 7 Å (dashed line), 6 Å (solid line), all with 0.15 subtraction of the mean intensity before French–Wilson correction of negative densities, and 6 Å (grey line) without any subtraction. (B) Detail of the 6 Å phasing map. (C) corresponding detail of the phase extended map at 2 Å resolution. Both maps are displayed at the same contour level.

The *Solomon* algorithm extended phases in 16 steps to 1.9 Å, using low-resolution phases of an X-ray density map of 1AUP for solvent flipping (40 s). Each phase extension step took 20 cycles, where in every fifth cycle the solvent mask was updated and in the subsequent cycle 40% of the protein density was truncated. The success of phase extension depended on the initial resolution of the phasing map, and phases of a 6 Å map were comfortably extended to 2.2 Å resolution. Remarkably, using these phase extension parameters, adding phase information with resolutions better than 6 Å did not further improve phase quality.

## 4. Discussion

Our benchmark shows that continuous electron-diffraction patterns can be oriented in three dimensions without corresponding real-space images or an assumed crystal lattice. *Diffractograph* preserves uncertainty in pairwise intersections: a match remains a probability distribution, including the possibility of a false match, until it is assessed against the geometry supported by other frames. This lets multiple orientation models be tested and refined without forcing the dataset into a single global solution.

The graph of frames serves as scaffolding for building those models, rather than as the reconstruction itself. *Diffractograph* explores enough of the graph to identify mutually consistent local models, then uses their refined orientations to guide further mergers. This differs from methods that first assign a single global set of pairwise relationships. It also allows incompatible models to remain separate, which may be useful for heterogeneous samples.

The method combines intensity correlations along predicted diffraction intersections with a probabilistic treatment of angular matches. Unlike approaches that select a common line or arc and then assess its reliability, *Diffractograph* retains the angular likelihood, its covariance and the probability of a false match. Evidence from several edges can therefore constrain a model even when individual orientations are ambiguous or weakly supported.

The benchmark includes Ewald-sphere curvature and dynamical electron scattering, although the orientation search treats each pattern as a planar section. The recovered orientations indicate that this approximation was adequate for the tested resolution range. A 3D reconstruction can then use the frame orientations to place the measured samples at their calibrated Ewald-sphere coordinates. The current results establish performance on this benchmark, which was also used during method development; they do not establish performance across experimental conditions. Further evaluation should vary signal-to-noise ratio, background, detector gaps and masking, beam-centre accuracy, angular coverage and graph connectivity, and should include heterogeneous particles and frames without useful particle signal.

The reconstructed 3D intensities could be phased by molecular replacement using a low-resolution map, followed by density modification and phase extension. This demonstrates that orientation recovery and intensity reconstruction provide useful input to downstream structure determination, rather than ending with a 3D intensity volume.

Together with the improved information efficiency reported for narrow-beam 4D-para-STEM [1], these results support a route towards 3D single-molecule electron diffraction: select frames with convincing C2-symmetric anisotropy, orient a stringent initial subset, then extend supported models with less decisive frames. The resulting orientations enable 3D intensity reconstruction; phase recovery can then use a low-resolution map and density modification.

## 5. Methods

### 5.1. Synthetic electron-diffraction benchmark

The diffraction patterns of glutamate dehydrogenase (PDB id: 1AUP) were created by multislice simulation as implemented in the abTEM package [20]. A combined Z-Y-Z Euler rotation matrix, defined by the angles φ, θ, and ψ, was applied. Prior to saving, these specific rotation angles were dynamically appended to the output file names to validate the resulting structural orientations.

The absolute square of the discrete Fourier transform of the exit wave, yields the intensity distribution in diffraction plane. The diffraction patterns contain information of up to 20 mrad and the multislice simulation was done using 200 keV.

The resulting benchmark dataset had 1741 diffraction patterns, stored in TIFF format, each corresponding to a specific rotation angle. The patterns included Ewald-sphere curvature and dynamical electron scattering and were generated independently of *Diffractograph*. The virtual detector had unit pixel size and a sample-to-detector distance of 4650 pixel units.

## 5.2. Software and computational environment

*Diffractograph* was implemented in ISO C++23. OpenAI Codex (OpenAI) was used to assist with software development and debugging. The authors reviewed and tested the resulting code and take responsibility for the software and results. Angular Fourier transforms were calculated using FFTW3 [21], and independent edge calculations, model operations and refinements were executed concurrently.

The reported reconstruction used *Diffractograph*, compiled with clang 21.0.0 under full optimization (-O3) and checking compatibility with the ISO standard using the -pedantic option. It was run on M1 Ultra, with 20 CPU cores, 64 GB RAM and MacOS Tahoe 26.5.2, using all CPU cores.

*Diffractograph*’s stages (orient, refine, complete, C1, 3D and phase) are separate executables that output JSON reconstruction reports containing provenance and results that are passed on to the next step.

## 6. Data and code availability

The *Diffractograph* source code and the synthetic 1741-frame benchmark data are available at https://github.com/senikm/diffractograph under the MIT license.

## Author contributions

J.P.A. conceived the probabilistic graph and model-growth method, implemented *Diffractograph*, performed the reconstruction analysis, and drafted the manuscript. S.M. developed and evaluated earlier diffraction orientation graph approaches, generated the synthetic benchmark dataset and its ground-truth orientations, and contributed to validation and revision of the manuscript. Both authors interpreted the results and approved the final manuscript.

